## Supplemental Figures for "Adult-onset CNS sulfatide deficiency causes sex-dependent metabolic disruption in aging"

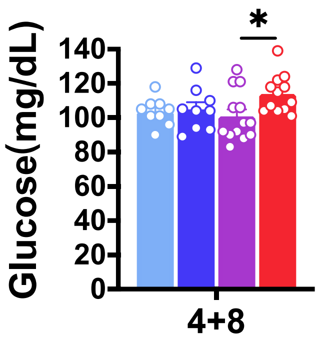


Fig. S1. Blood glucose levels measured without fasting at 8 months post tamoxefin injection under normal diet. n=9-14. Multiple t-Test.

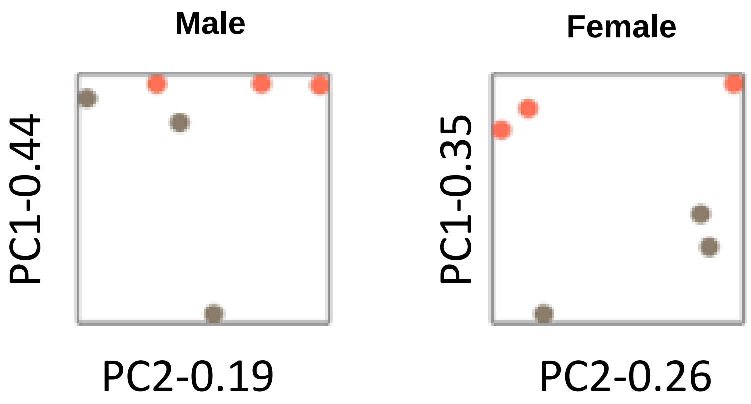


Fig. S2. PCA showing that female CST Cre+ cluster in a more distinct manner versus Cre- female mice relative to male CST Cre+ and Cre- mice

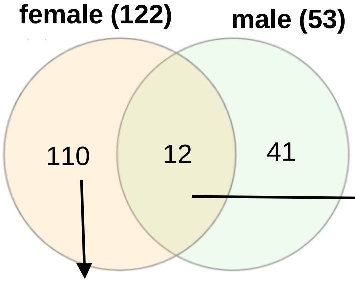


| | | | |
|---|---|---|---|
| Bcl2a1a | Ccl3 | Ccl4 | Cd86 |
| Ch25h | Gal3st1 | lag3 | Mmp12 |
| Opalin | Ptpn6 | Serina3n | Timp1 |

| | | | | | | | | | |
|---|---|---|---|---|---|---|---|---|---|
| Apoe | Atf3 | Axl | Bag4 | Bcl10 | Blnk | Bok | C1qa | C1qb | C1qc |
| C3 | C3ar1 | C4a | C5ar1 | Cables1 | Casp7 | Ccl2 | Ccl5 | Ccl7 | Cd109 |
| C4 | Cd33 | Cd68 | Cd72 | Cd84 | Clec7a | Cotl1 | Csf1 | Csf1r | Csf3r |
| Cst7 | Ctss | Cx3cr1 | Dock2 | Erbb3 | F3 | Fabp5 | Fbln5 | Fcer1g | Fcgr1 |
| Fcgr2b | Fcgr3 | Fcrls | Fgd2 | Gadd45a | Gadd45g | Gpr183 | Gpr34 | Grn | Hmox1 |
| Hsd11b1 | Hspb1 | Icam2 | Ifi30 | Igsf6 | Il2rg | Il3ra | Inpp5d | Irak2 | Irf8 |
| Itgax | Itgb5 | Jag1 | Lilrb4a | Ly9 | Lyn | Mafb | Man2b1 | Mbd3 | Mcm2 |
| Mef2c | Mpeg1 | Mr1 | Msn | Mvp | Myc | Ncf1 | Nfkb2 | Osmr | Pdpn |
| Pla2g4a | Plcg2 | Pros1 | Psmb8 | Rad51c | Relb | Rsad2 | S100a10 | S1pr3 | Siglecf |
| Slamf9 | Slc10a6 | Slco2b1 | Sox9 | Spp1 | Tcirg1 | Tgfb1 | Tgfbr1 | Tgm1 | Tlr2 |
| Tlr7 | Tm4sf1 | Tmem204 | Tnfrsf1a | Traf2 | Trem2 | Trp53 | Tyrobp | Vav1 | Vim |

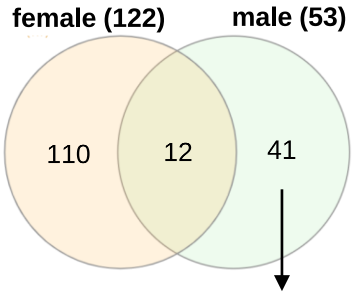


| | | | | | | | | | |
|---|---|---|---|---|---|---|---|---|---|
| Atg3 | Atg5 | Atm | Brd2 | Casp8 | Cd69 | Cd70 | Creb1 | Csk | Ctse |
| E2f1 | Esam | Fcrla | Fdxr | Foxp3 | Gpr84 | Igf1 | Il21r | Irf2 | Irf7 |
| Islr2 | Itga6 | Kcnk13 | Kdm4a | Klrk1 | Kmt2c | Lig1 | Ly6a | Myct1 | Pnoc |
| Ptx3 | Rab6b | Ripk1 | Setd7 | Sin3a | Slamf8 | Snca | Tmem119 | Tnfrsf11b | Tnfsf8 |
| Vps4a | | | | | | | | | |

Fig. S3. DEGs lists in Venn diagrams from female or male CST $Cre^{+}$ vs. CST $Cre^{-}$.
